# Behavioural effects of temperature on ectothermic animals: unifying thermal physiology and behavioural plasticity

**DOI:** 10.1101/056051

**Authors:** Paul K. Abram, Guy Boivin, Joffrey Moiroux, Jacques Brodeur

## Abstract

Temperature imposes significant constraints on ectothermic animals, and these organisms have evolved numerous adaptations to respond to these constraints. While the impacts of temperature on the physiology of ectotherms have been extensively studied, there are currently no frameworks available that outline the multiple and often simultaneous pathways by which temperature can affect behaviour. Drawing from the literature on insects, we propose a unified framework that should apply to all ectothermic animals, generalizing temperature's behavioural effects into (1) Kinetic effects, resulting from temperature's bottom-up constraining influence on metabolism and neurophysiology over a range of timescales (from short-to long-term), and (2) Integrated effects, where the top-down integration of thermal information intentionally initiates or modifies a behaviour (behavioural thermoregulation, thermal orientation, thermosensory behavioural adjustments). We discuss the difficulty in distinguishing adaptive behavioural changes due to temperature from behavioural changes that are the products of constraints, and propose two complementary approaches to help make this distinction and class behaviours according to our framework: (i) behavioural kinetic null modeling and (ii) behavioural ecology experiments using temperature-insensitive mutants. Our framework should help to guide future research on the complex relationship between temperature and behaviour in ectothermic animals.

## 1. Introduction

Temperature influences all levels of biological organization. As temperature increases, the higher kinetic energy of biochemical reactions speeds up the rate of metabolic processes, scaling up to affect the physiology and behaviour of individual organisms (Angilletta 2009) and ecosystem-level processes (Brown et al. 2004). This thermal rate dependency of physiology and behaviour is the most predictable for ectothermic organisms, which, in contrast to endotherms, do not typically maintain a constant body temperature through homeostatic processes.

While the effects of temperature on ectothermic organisms have long been recognized, research in this area has recently intensified (e.g., Deutsch et al. 2008; Dell et al. 2011; Dell et al. 2014; Huey et al. 2012; Gilbert et al. 2014; Sunday et al. 2014; Vasseur et al. 2014; Woods et al. 2014). In part, this renewed interest has been spurred by a need to understand the response of individuals and ecosystems to recent and incipient climate change, wherein average global temperatures are increasing, and extreme temperature events are becoming more commonplace (Stocker et al. 2013). Due to the extensive effects these changes are having on ecosystems, it is critical to understand the basic details of how species and ecosystems cope with temperature.

In parallel, seminal studies by Gillooly et al. (2001) and Brown et al. (2004), among others, have proposed that the rates of many large-scale ecological phenomena (e.g. population growth rates, trophic interactions, biomass production) can be in large part predicted by metabolic rates of individual organisms that increase exponentially with increasing temperature, as predicted by first-principles enzyme kinetics (i.e., the Metabolic Theory of Ecology). While these macroecological approaches have been very influential and lend hope to the prospect of understanding large-scale processes from simple mathematical principles, they have a number of key mechanistic and predictive deficiencies that have received substantial criticism (Cyr and Walker 2004; Clarke 2006; O'Connor et al. 2007; Irlich et al. 2009). From a behavioural ecology perspective, one important critique is that this global view tends to ignore much of what is happening at intermediate levels of biological organization. For example, many ectothermic organisms have the capacity to adapt their physiology and behaviour to modify or better tolerate their thermal environment. That is, temperature not only modifies how (e.g. how rapidly) behaviours are performed, but can determine whether or not a given behaviour is expressed or which one of a possible set of behaviours is performed in a given situation (i.e., behavioural “decisions”; *sensu* Dell et al. 2011). Some of these behavioural changes are likely mediated by direct perception of temperature: an exciting collection of recent studies in *Drosophila* fruit flies have begun to investigate the neurological mechanisms involved in the reception and transmission of thermal information and its control over thermoregulatory behaviour (e.g. Hamada et al. 2008; Gallio et al. 2011; Frank et al.2015; Liu et al. 2015). However, the potential of this “top-down” integration of thermal information to drive other forms of adaptive behavioural plasticity, and its interaction with more straightforward metabolic effects of temperature, have not been given their due consideration by physiologists or behavioural ecologists.

In this paper, we (1) synthesize the empirical research on the effect of temperature on the behaviour of ectotherms, focusing on insects, and (2) develop a framework to categorize these behavioural responses, considering the often simultaneous influences of constraints imposed by temperature and adaptive behavioural plasticity in response to temperature perception. Although based on insect studies, this framework applies to all ectothermic animals. Our synthesis highlights that a lack of organized understanding of thermal effects on behaviour may have lead to a rarity of hypothesis-based research concerning temperature-induced adaptive behavioural modifications. Furthermore, there is currently relatively limited evidence to support the notion that individuals' decisions take thermal information into direct account. We discuss how these impediments could potentially be overcome. Our framework will hopefully serve as a guideline for future research into the effects of temperature on the behaviour of ectotherms, and will contribute to a more complete understanding of thermal biology across all levels of biological organization.

## 2. Definitions and Scope

In order to categorize the effects of temperature on ectothermic animal behaviour, what constitutes behaviour must first be defined. Here, we employ the general definition formulated by Levitis et al. (2009):

> *“Behaviour is the internally coordinated responses (actions or inactions) of whole living organisms (individuals or groups) to internal and/or external stimuli, excluding responses more easily understood as developmental changes.”*

The key component of this definition is the concept of an “internally coordinated response by the whole organism”, which distinguishes underlying physiological mechanisms behind behaviours-often at the level of a cell, tissue, or organ-from the whole-organism response (behaviour) that occurs when these physiological responses are coordinated (e.g., via the neural and endocrine systems). We conducted our literature search while adhering to this definition, but also accepted that grey areas often arise when attempting to classify organismal responses across the physiology-behaviour continuum (see Levitis et al. 2009 for an extensive discussion).

Because the literature describing temperature's effects on ectothermic animal behaviour and physiology is vast, we limited the scope of our literature search to insects. This group of ectotherms is well-studied in terms of their extremely diverse behaviours under different environmental conditions, the underlying physiological, cellular, and molecular mechanisms of these behaviours, and the understanding of behaviour's connection to ecological processes. We further limited our scope to thermal modifications of behaviour occurring within insects' non-stressful range of temperatures. In traditional thermal performance curves (see **section 3.1**), this could be viewed as the exponential, increasing portion of the curve from the critical thermal minimum to the optimal temperature (Angilletta 2009). Therefore, we did not consider behavioural changes in response to heat or cold shock, fluctuating temperatures, hardening, or behaviours resulting from permanent or semi-permanent nervous or tissue damage caused by extreme low or high temperatures (see Chown and Nicolson 2004 pp. 115-150; Hance et al. 2007; Bowler and Terblanche 2008]; Colinet et al. 2015 for reviews on these subjects). Furthermore, we did not include the processes of diapause or overwintering, which, although some times referred to as behaviours, we considered physiological/developmental states due to the relatively long time scale over which they occur. However, component behaviours involved in preparation or exit from diapause/overwintering would in principle fall within the scope of our discussion.

By identifying patterns in the literature, based on underlying mechanisms and behavioural outputs, we constructed a general framework to permit the categorization of all possible behavioural responses to temperature in ectothermic animals. We distinguished two main components of behavioural responses to temperature: “Kinetic” and “Integrated” effects, and further subdivided them based on mechanisms and outcomes. We will discuss each in turn, providing examples from the literature that can be classified according to this new framework.

## 3. Kinetic effects

Many components of temperature-induced behavioural changes in animals ultimately result from a constraint: the amount of kinetic energy present in physiological systems. Higher kinetic energy (due to higher temperature) increases the rate of conformational change of proteins (including enzymes), as well as the proportion of reactants that reach the activation energy required for biochemical reactions to occur (reviewed in Fields 2001; Hochachka and Somero 2002). Higher temperature also increases the fluidity of cell membranes, affecting the transport of materials in and out of cells (e.g., Hazel and Williams 1990; Hazel 1995). These biochemical effects propagate through biological levels of organization in a bottom-up fashion to affect the speed of cell, organ, system (muscular, nervous, digestive), and finally whole-organism behavioural processes (Fig. 1). These “Kinetic effects” can be expressed over a continuum of timeframes, which in subsequent sections we will subdivide into short-term (within a matter of seconds or minutes), medium-term (over hours, days, or weeks within the same ontogenetic stage), and long-term (across ontogenetic stages) effects.

**Fig. 1.**
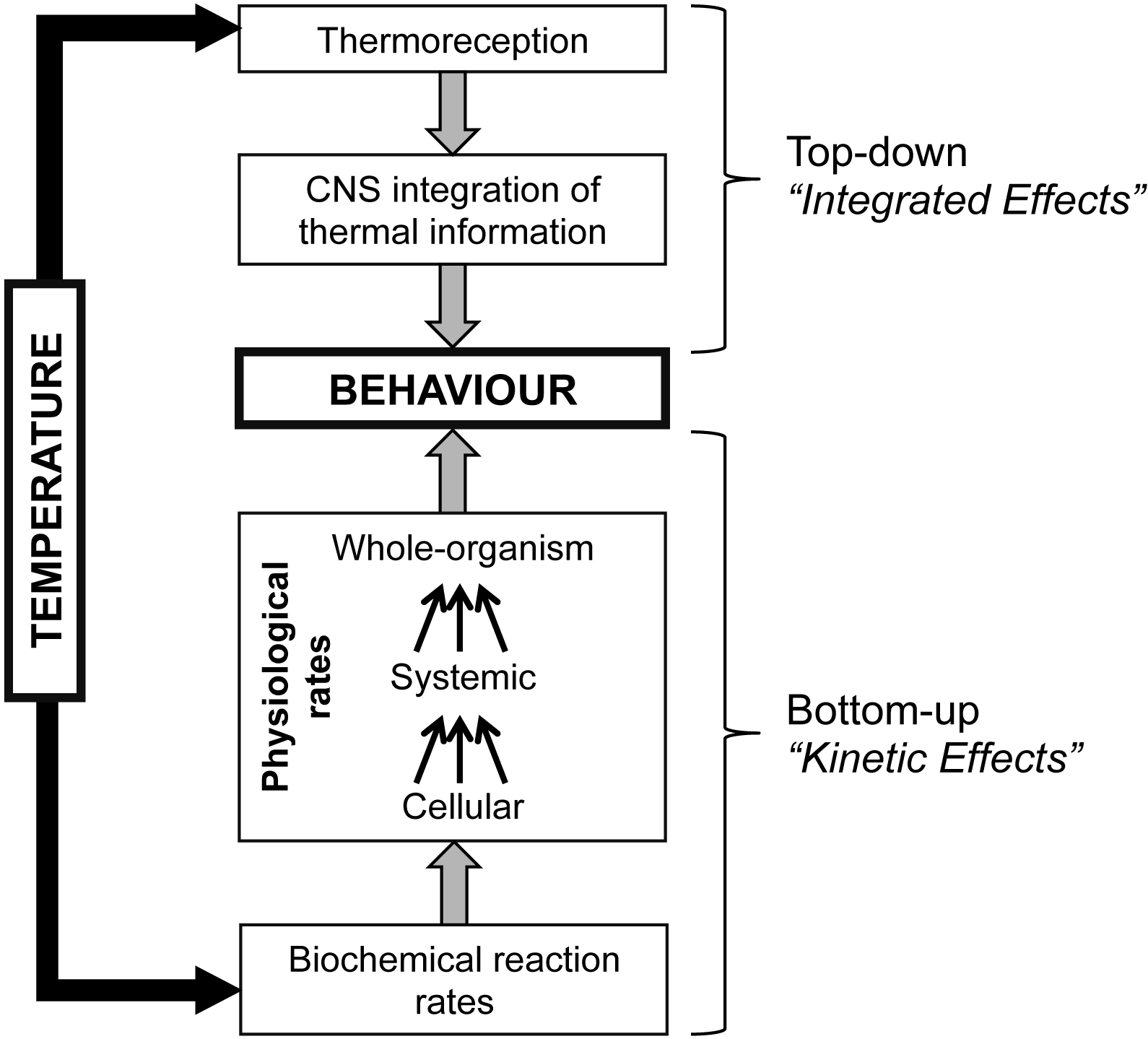
Kinetic and Integrated effects of temperature on the behaviour of ectothermic animals. Kinetic effects (bottom-up) ultimately result from changes in biochemical reaction rates, which scale up via multiple cellular and systemic physiological processes to influence whole-organism processes (i.e. behaviour). Integrated effects (top-down) result from thermoreception and integration of thermal information by the central nervous system (CNS), which then modulates behavioural outputs. Both Kinetic and Integrated effects act in concert to determine an animal's behavioural phenotype.

### 3.1 Short-term Kinetic effects

The best-characterized effect of temperature on ectothermic animal behaviour is an increasing rate at which behaviours are performed with increasing temperature. For example, ambient temperature affects the incidence and rate of locomotor (e.g. walking speed, flight performance), reproductive (mating, oviposition), and foraging (search, attack) behaviours (Table 1). We classify these changes, which can typically be observed within seconds or minutes of a temperature change, as short-term Kinetic effects (Fig. 2).

**Fig. 2.**
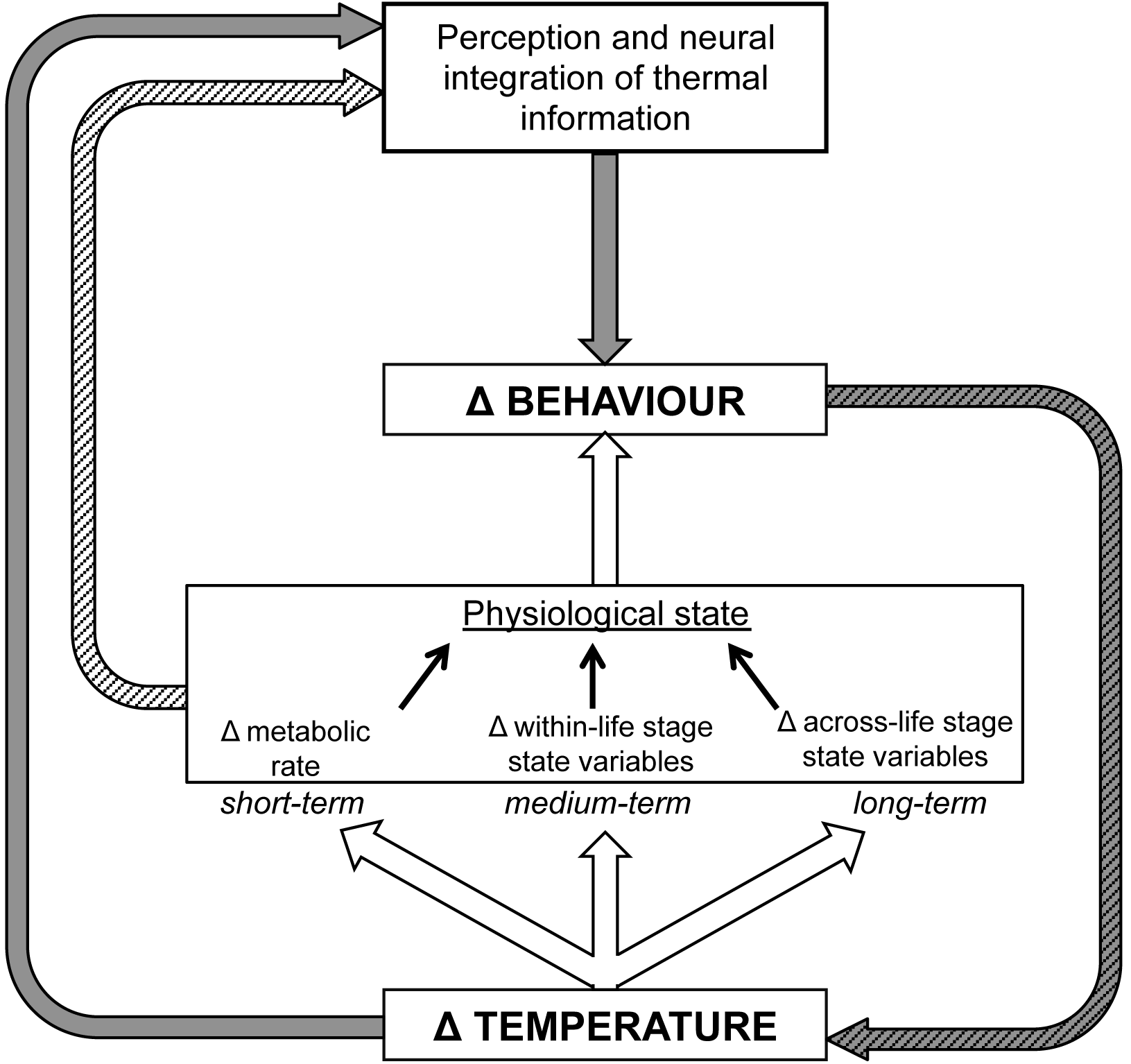
A general schematic for how temperature changes (Δ) elicit behavioural changes in ectothermic animals. Kinetic effects (white arrows) act on aspects of an animal's physiological state and resulting behaviour across a range of timeframes, from short-term (causing changes in metabolic rate on the range of seconds to minutes) to long-term (causing differences in state variables across life stages, e.g. by influencing resource allocation during development). Integrated effects (grey arrows), resulting from temperature perception, intentionally modulate behaviour after being integrated by the central nervous system (thermosensory behavioural adjustments, thermal orientation). In some cases, this behavioural change involves the animal changing its own thermal environment (behavioural thermoregulation; hatched grey arrow). How thermal information is perceived and/or integrated, and resulting downstream behavioural changes due to Integrated effects, can also be modulated by thermally-dependent physiological state (white hatched arrow).

**Table 1.**
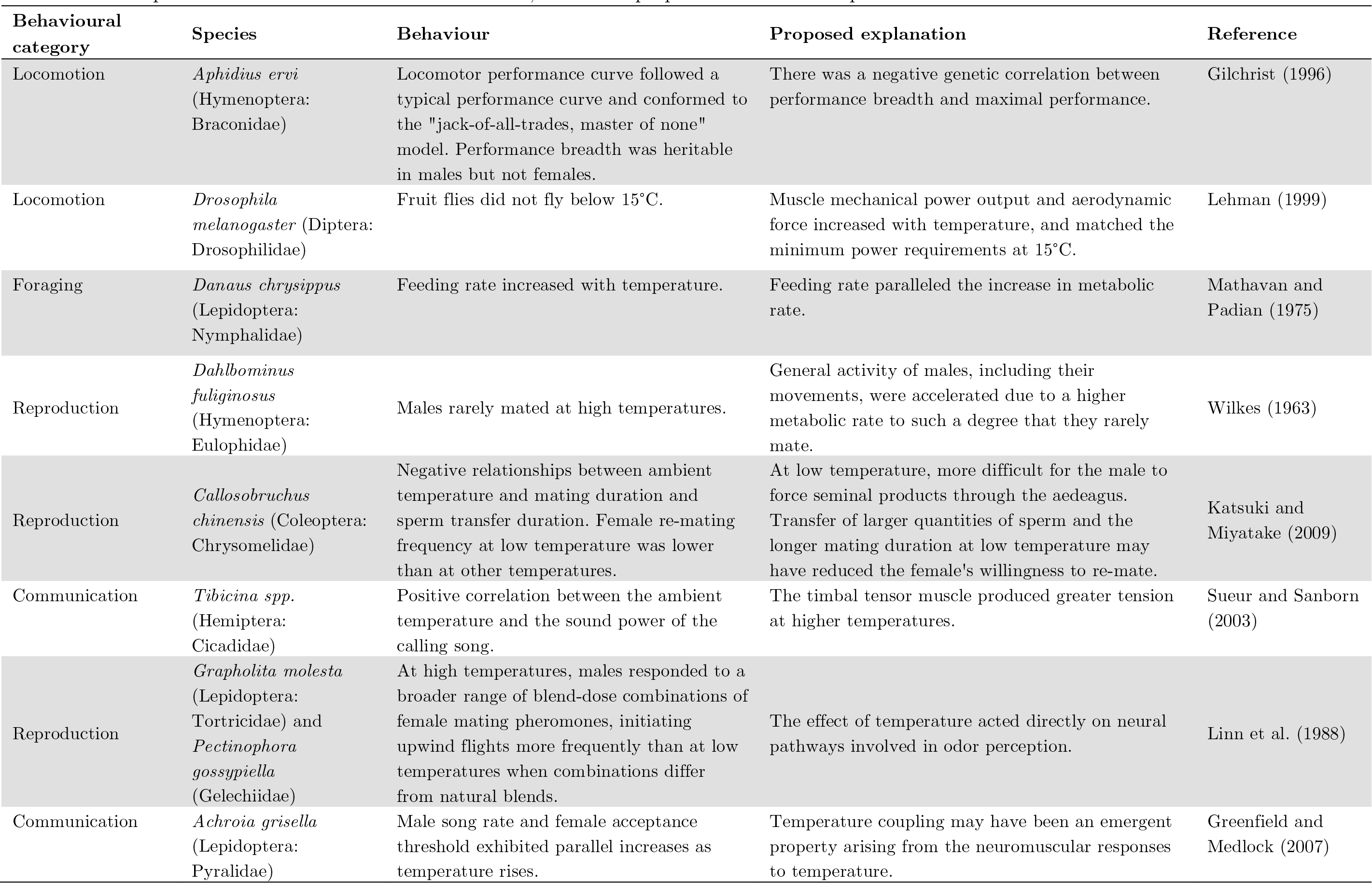
Examples of short-term Kinetic effects in insects, with their proposed mechanistic explanations

For the purposes of our framework, short-term Kinetic effects can invariably be described by “thermal performance curves” (Huey and Stevenson 1979), the application of which extends to any biological process whose rate is affected by temperature (e.g. development, digestion, egg production). Thermal performance curves are also often applied to behavioural measures of performance, which are also strongly dependent on metabolic rate (Dell et al. 2011). Above a certain critical thermal minimum, behaviours tend to be performed more rapidly as temperature increases up to an optimum temperature, their rate then decreasing up to a critical thermal maximum after which the behaviour can no longer be performed. There are a variety of approaches for characterizing and modeling thermal performance curves, which are discussed elsewhere (Angilletta 2006; Angilletta 2009; Damos et al. 2012; Regnière et al. 2012).

### 3.2 Medium-term Kinetic effects

Some of the behavioural effects of temperature on ectothermic animals occur after the original temperature exposure (Fig. 2), and thus cannot be described in real-time by thermal performance curves. For example, within insect life stages temperature exposure can change subsequent behavioural processes via its effects on various metabolism-dependent aspects of physiological state, which themselves have effects on the fitness payoffs of different behaviours or alternative variations on a single behaviour (Table 2). Thus, medium-term Kinetic effects can broadly be considered as thermal constraints acting on the underlying state-dependency of adaptive behavioural strategies. For example, many aspects of insect behaviour are sensitive to egg load (Minkenberg et al. 1992), and the rate at which females synthesize eggs is temperature-dependent (e.g., Berger et al. 2008). Therefore, behavioural variables such as searching intensity, clutch size, and host/prey handling time can be indirectly influenced by temperature, via its effect on egg load (Rosenheim and Rosen 1991; Berger et al. 2008; Table 2). Medium-term effects of temperature also include changes in insect behaviour due to the thermal dependency of state variables external to the individual animal itself. For example, temperature can alter the frequency of pauses during nesting behaviour by changing the developmental rate of offspring (Weissel et al. 2006), and modify oviposition behaviour by altering the composition, quantity, and persistence of chemical signals deposited by conspecific competitors (Sentis et al. 2015a) (Table 2). These kinds of temporally offset, indirect effects of temperatures on state variables and their consequences for animal behaviour remain relatively poorly studied compared to more straightforward short-term effects of temperature, but are likely to explain a considerable amount of behavioural variation in natural populations of ectothermic animals. Finally, a better-knownmedium-term effect of temperature on behaviour is known as reversible acclimation (*sensu* Angilletta 2009, p. 126), where exposure to a given temperature regime modifies the subsequent thermal sensitivity of behaviour (i.e., the shape of the thermal performance curve) within the same life stage, including those involved in foraging and reproduction (Sentis et al. 2015b; reviewed in Angilletta 2009; Chown and Nicholson 2004).

**Table 2.**
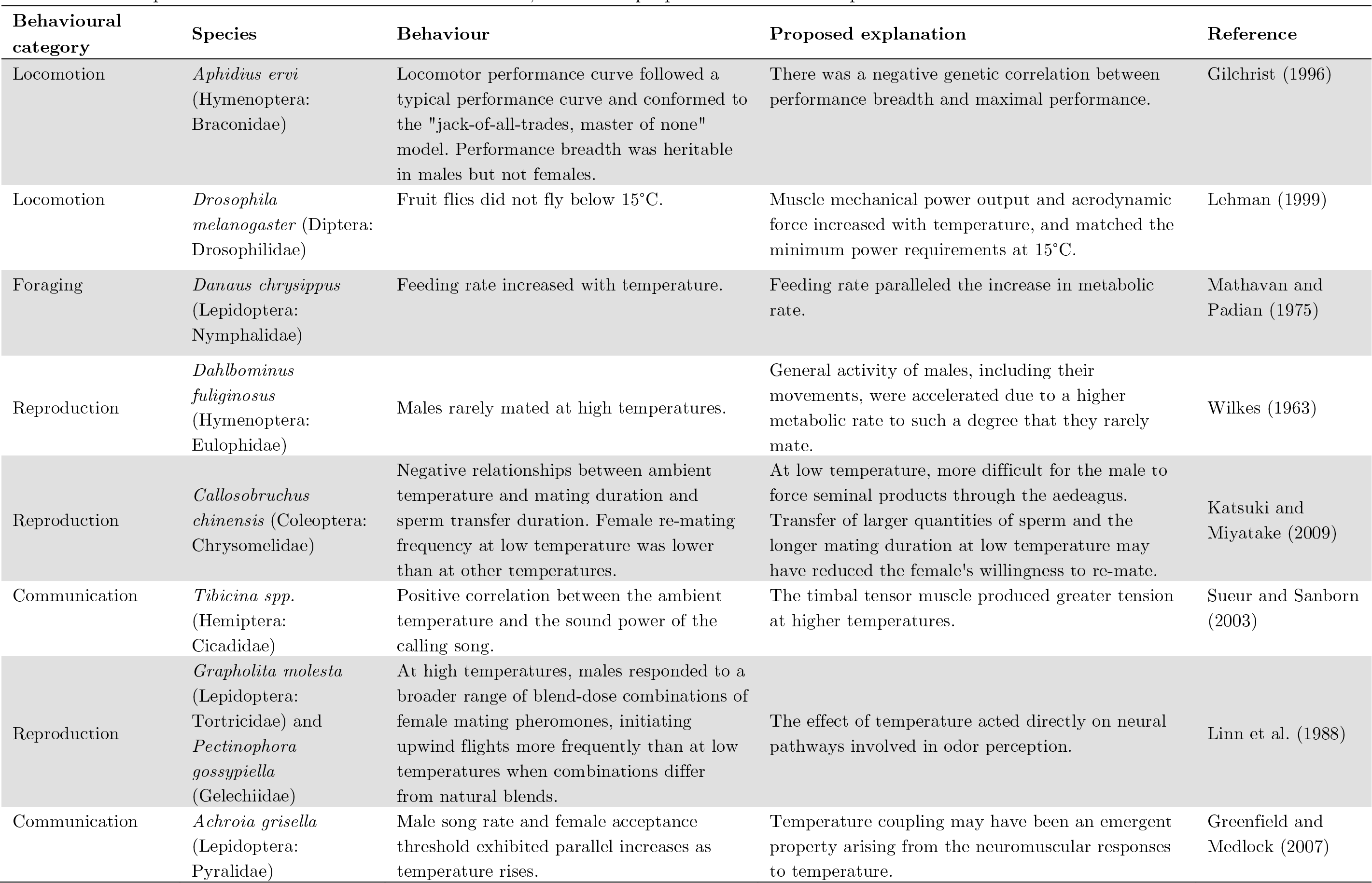
Examples of medium-term Kinetic effects in insects, with their proposed mechanistic explanations.

### 3.3 Long-term Kinetic effects

In addition to short- and medium-term effects, temperature's Kinetic effects on ectothermic animal behaviour can occur over longer time scales, acting across ontogenetic stages to shape behavioural phenotypes. Although the mechanistic bases of these changes remain controversial (e.g., Angilletta and Dunham 2003; Forster and Hirst 2012; Ghosh et al. 2013), developmental temperature does affect the relative allocation of developmental resources to different functions, resulting in phenotypic trait variation that can be described by thermal reaction norms (Angilletta et al. 2003; Kingsolver et al. 2004). In contrast to thermal performance curves, the phenotypic traits described by thermal reaction norms are typically fixed and irreversible for an individual within an ontogenetic stage. When the thermal sensitivity of performance in one ontogenetic stage is affected by the temperature experienced during the preceding stage, these effects are referred to as “developmental” or “irreversible” acclimation (Angilletta 2009; Cavieres et al. 2016), although it has been argued that use of the term acclimation is problematic because these changes are actually responses of developmental plasticity to temperature (Wilson and Franklin 2002). In addition to changes in thermal sensitivity, variation in other phenotypic traits due to developmental temperature can have a wide range of consequences for behaviour including locomotion, foraging, and mating (Table 3). For instance, in about three quarters of insects species, individuals that develop at lower temperatures tend to be larger than those that develop at higher temperatures, a phenomenon termed the “temperature-size rule” (Atkinson 1994). Because size itself has large effects on behavioural strategies, the temperature-size rule is a common way for developmental temperature to have long-term consequences for behavioural strategies. For example, insect parasitoids that are smaller as a result of developing at higher temperatures may have altered fecundity, basal metabolic rates, and life expectancy, changing the efficiency with which they interact with hosts and exploit host patches (Colinet et al. 2007; Le Lann et al. 2011). Due to their smaller size, warm-developing parasitoids tend to be less capable of subduing larger hosts (Le Lann et al. 2011; Wu et al. 2011; Moiroux et al. 2015) and may adopt more risk-prone behaviour (Moiroux et al. 2015), possibly representing adaptive behavioural plasticity in response to developmental temperature-induced size variation.

**Table 3.** Examples of long-term Kinetic effects in insects, with proposed mechanistic explanations

| Behavioural category | Species | Behaviour | Proposed explanation | Reference |
| --- | --- | --- | --- | --- |
| Foraging | *Aphidius rhopalosiphi*<br>(Hymenoptera: Braconidae) | Oviposition rate was higher, and patch residence time and host handling time were shorter in females reared at low temperature than females reared at high temperature when foraging at a common temperature. | A higher metabolic rate and greater size when reared at low temperatures meant that females were more active, moved faster and parasitized more hosts. | Le Lann et al. (2011) |
| Locomotion | *Drosophila melanogaster*<br>(Diptera: Drosophilidae) | Cold-reared fruit flies failed to perform a take-off flight less frequently than warm-reared flies. | The improved ability to fly at low temperatures was associated with a dramatic increase in wing area and length. | Frazier et al. (2008) |
| Reproduction | *Aphidius colemani*<br>(Hymenoptera: Braconidae) | Females that developed at low and high temperatures laid more eggs than females developing at intermediate temperatures. | The decrease in oviposition rate at high temperature may be the result of a decrease in fat reserves when development time was shorter. At low temperatures, standard thermal rate-dependency causes lower oviposition rates. | Colinet et al. (2007) |
| Reproduction | *Drosophila melanogaster*<br>(Diptera: Drosophilidae) | Males reared at 25°C had greater territorial success than males reared at 18°C and males whose parents developed at 25°C were more successful than were males whose parents developed at 18°C. | Fruit flies raised at 25°C, even though phenotypically small, are physiologically more vigorous than flies reared at 18°C. | Zamudio et al. (1995) |
| Learning | *Apis mellifera*<br>(Hymenoptera: Apidae) | Bees reared at 35°C had a better short-term memory than bees reared at 32°C, while rearing temperature had no significant effect on the long-term learning and memory abilities of workers. | Temperature had no effect on fluctuating asymmetry. Consequences of rearing temperatures are thus subtle neural deficiencies affecting short-term memory rather than physical abnormalities. | Jones et al. (2005) |
| Foraging | *Apis mellifera*<br>(Hymenoptera: Apidae) | Honeybees raised at high temperatures showed an increased probability to dance, foraged earlier in life, and were more often engaged in removal of dead colony members. | Low pupal developmental temperatures lead to reduced juvenile hormone (JH) biosynthesis rates in adult bees, causing delayed behavioural development and hence a later onset of foraging. Further, developmental temperature may have interfered with JH metabolism and/or octopamine levels in the bee's brain, affecting foraging preferences. | Becher et al. (2009) |

Beyond the temperature-size rule, the metabolic compensation hypothesis is a possible source of long-term Kinetic effects of temperature on insect behaviour. This hypothesis, which has been rarely investigated-but has been experimentally confirmed in *Drosophila* (Berrigan and Partridge 1997) and parasitoids (Le Lann et al. 2011)-posits that cold-developed individuals tend to have a higher metabolic rate than warm-developed individuals when placed at a common temperature, probably because of a higher production of mitochondria and associated enzymes at low temperatures (Hochachka and Somero 2002). Le Lann et al. (2011) observed that this higher metabolic rate induced by low developmental temperatures resulted in decreased adult longevity, but was counterbalanced by a higher egg load in an aphid parasitoid. These authors proposed that this change in resource allocation may explain why cold-developed females exploited patches more intensively than warm-developed females, and suggested that metabolic rate should be considered more frequently when investigating the influence of developmental temperature on behavioural decisions.

## 4. Integrated effects

So far we have considered effects of temperature on behaviour that ultimately result from bottom-up effects of the amount of kinetic energy present in physiological systems. Now we will discuss how organisms can respond to thermal information received by thermosensory organs, which is then integrated in the central nervous system (CNS) and used to intentionally modulate behaviours in a top-down fashion (Fig. 1). We term these behavioural changes “Integrated effects”.

Insects have the capacity to sense temperature and modify their behaviour as a result. External thermosensory organs (e.g. sensilla) have been identified for many insects, often residing on antennae but also found on other areas of the body such as the thorax, head, legs, and ovipositor (Altner and Loftus 1985; Brown and Anderson 1998; Zars 2001). Internal thermosensory anterior cell neurons located in the brain have also been identified in *Drosophila* (Hamada et al. 2008). Recent detailed studies of *Drosophila* thermotaxis demonstrate that both external thermosensing organs and internal thermosensory neurons play a key role in behavioural thermoregulation. Temperature sensation by these structures is mediated by an array of Transient Receptor Protein (TRP) cation channels that are activated by temperatures above or below the flies' preferred temperature (∼25°C) (Sayeed and Benzer 1996; Zars 2001; Hamada et al. 2008; Gallio et al. 2011; Shen et al. 2011; Fowler and Montell 2012). In *Drosophila* adults, most peripheral thermosensory neurons are located in the antennae. A subset of these neurons are excited by cooling and inhibited by warming (‘cold-sensitive’), in opposition to ‘warm-sensitive’ thermoreceptors, which are excited by warming and inhibited by cooling (Hamada et al. 2008; Gallio et al. 2011). These sensory inputs produce a thermal 'map’ in the *Drosophila* brain (Gallio et al. 2011) which is further processed by projection neurons in the proximal antennal protocerebrum and passed on to several other brain regions implicated in sensory reception and behavioural modulation (Frank et al. 2015; Liu et al. 2015). These sophisticated systems of thermoreceptors and neural networks provide fine-scale thermal information that is required for flies to undertake normal thermoregulatory behaviour (Frank et al. 2015; Gallio et al. 2011). The neurological basis of temperature sensation has also been explored in leaf-cutting ants, which utilize neurologically integrated thermal information as an orientation cue (Kleineidam et al. 2007; Ruchty et al. 2010a; Ruchty et al. 2010b; Nagel and Kleineidam 2015). Taken together, these mechanistic studies remind us that the effects of temperature on insect behaviour are not purely metabolic: these animals can intentionally modify their behaviour as a result of temperature perception. Studies on the effects of integrated thermal information for insect behaviour and ecology are limited to very few study organisms, have mostly been conducted in the laboratory, and have not been investigated outside of the context of thermotaxis and thermal orientation-but there is no reason why integrated thermal information could not be used to modulate other behavioural processes. In the three following sections, we will class Integrated effects of temperature on behaviour depending on whether they result in a change in the temperature experienced by the organism (behavioural thermoregulation), are used by the organism as a navigational tool (thermal orientation), or rather adjusted decision-making that compensates for or takes advantage of the temperature change (thermosensory behavioural adjustments).

### 4.1 Behavioural thermoregulation

Ectothermic animals are not completely constrained by the Kinetic effects imposed by ambient temperatures in a given location. While ectotherms cannot typically adjust their body temperatures through endogenous homeostatic control (as in endotherms), the fact that natural environments are thermally heterogeneous allows the modification of body temperature via behavioural thermoregulation (May 1979; Woods et al. 2014). The mechanistic studies of *Drosophila* thermoreception described above suggest that these behaviours are likely to be largely mediated by the top-down effects of integrated thermal information. Thus, in the case of thermoregulation, the ambient temperature experienced by an animal causes a behavioural response, which in turn changes ambient temperature, which can subsequently affect physiological functions via Kinetic effects (Fig. 2).

Thermoregulatory behaviours in insects are diverse and the literature describing them extensive, but they can be grouped into the broad categories of (i) microhabitat selection, (ii) basking, (iii) modification of temporal activity cycles and (iv) endothermy (see May 1979 for an early review). Insects use a given strategy, or a combination of several, to avoid aversive temperatures and maintain a body temperature that optimizes such diverse functions as reproduction (Hedrick et al. 2002; Spence et al. 1980; Stahlschmidt and Adamo 2013), growth and development (Clissold et al. 2013; Coggan et al. 2011), locomotion (Dorsett 1962; Kingsolver 1985; Whitman 1988; Dreisig 1981), and defense against parasitic infection (ie. ‘behavioural fever’) (Boorstein and Ewald 1987; Elliot et al. 2002; Ouedraogo et al. 2004). Some representative examples of thermoregulatory behaviours and their proposed functions are shown in Table 4.

Behavioural thermoregulation has a critical role in shaping ectotherms' evolutionary adaptations to environmental thermal variation. While behaviour is often seen as a driver of evolutionary change (e.g., Duckworth 2009; Zuk et al. 2014), Huey et al. (2003), building on longstanding hypotheses (Bogert 1949), elegantly demonstrated that behavioural thermoregulation might actually act as an inertial force for the evolution of adaptations to temperature variation (the “Bogert effect”). Because behavioural thermoregulation essentially functions to mitigate thermal variations experienced by individuals that do not thermoregulate, it would reduce the strength of selection on other temperature-dependent physiological or behavioural traits (Huey et al. 2003). In addition, behavioural thermoregulation could buffer many of the negative effects of climate change on animals, although this will depend on factors such as the degree of thermal specialization and the availability of opportunities for thermoregulation (Kearney et al. 2009; Huey and Tewksbury 2009; Huey et al. 2012; Sunday et al. 2014).

Behavioural thermoregulation strategies are dynamic, since the optimal temperature for a given individual can depend on the thermal optima of different aspects of physiological state (Huey 1991). For example, *Locusta migratoria* (L.) (Orthoptera: Acrididae) nymphs move to lower temperatures along a thermal gradient with increasing degrees of nutrient deprivation, which increases the efficiency of protein and carbohydrate assimilation at the cost of lower rates of growth and development (Coggan et al. 2011). These locusts can even use this dynamic thermoregulatory behaviour to optimize assimilation of specific nutrients, which varies with temperature (Clissold et al. 2013). Similarly, the preferred temperature of *Triatoma* bugs (Hemiptera: Reduviidae) along a gradient depends on nutritional status, with bugs preferring cooler temperatures when starved, and warmer temperatures following a blood meal (Guarneri et al. 2003; Minoli and Lazzari 2003). *Triatoma* thermopreference also has a strong circadian component, with bugs preferring warmer temperatures just before the onset of darkness; this cycle persists even in the absence of light/dark cues, suggesting the existence of an internal clock for thermal preferences (Minoli and Lazzari 2003). These examples, along with the others provided (Table 4), support the hypothesis that the behaviours involved in thermoregulation likely receive complex neural inputs integrating differenttypes of physiological information, including top-down Integrated thermal information (Gallio et al. 2011; Frank et al. 2015), as well as bottom-up information on state variables (e.g. nutritional status) that are subject to short-term and medium-term Kinetic effects of temperature.

**Table 4.** Examples of thermoregulatory behaviours in insects, with their proposed functions.

| Behavioural category | Species | Behaviour | Proposed function | Reference |
| --- | --- | --- | --- | --- |
| Microhabitat selection | *Gryllus integer*<br>(Orthoptera: Gryllidae) | Male crickets chose cracks in the ground at a temperature where the frequency and intensity of their calling songs were maximized. | Optimize attractive characteristics (e.g., chirp rate) of calling songs to mates. | Hedrick et al. (2002) |
| Microhabitat selection | *Locusta migratoria*<br>(Orthoptera: Acrididae) | Locust nymphs deprived of protein or carbohydrate and then fed a meal subsequently chose different temperatures. | Maximize the assimilation of the nutrient that addresses their nutritional imbalance. | Clissold et al. (2013) |
| Microhabitat selection | *Bombus terrestris*<br>(Hymenoptera: Apidae) | Bees parasitized by conopid flies stayed outside the nest at night and actively sought out cooler temperatures. | Retard the development of the parasitoid (prolonging the life of the bee) and reduce the parasitoid's developmental success. | Müller and Schmid-Hempel (1993) |
| Microhabitat selection | *Solenopsis invicta*<br>(Hymenoptera: Formicidae) | When ant colonies were well-fed, workers and brood were kept at high temperatures; when colony was food-limited, brood were still kept at high temperatures but workers were not associated with brood care seek lower temperatures. | Increase the longevity of workers not directly associated with brood care, while continuing to optimize brood development temperature, when resources are limited. | Porter and Tschinkel (1993) |
| Basking | *Pieris* spp. (Lepidoptera: Pieridae) | Butterflies changed orientation of their wings to reflect solar radiation onto their thorax. | Attain body temperatures necessary for flight. | Kingsolver (1985) |
| Basking | *Gerris* spp. (Hemiptera: Gerridae) | Adult water striders submerged themselves underwater when water temperature was higher than ambient air temperature. | Increase the rate of gonad maturation and egg production. | Spence et al. (1980) |
| Daily activity cycle | *Drosophila subobscura*<br>(Diptera: Drosophilidae) | Flies at warm locations were mostly active in the evening. | Avoid exposure of eggs to high daytime temperatures soon after oviposition. | Huey and Pascual (2009) |
| Microhabitat selection, basking, daily activity cycle | *Taeniopoda eques*<br>(Orthoptera: Romaleidae) | Grasshoppers (i) selected sites with large amounts of solar radiation and (ii) "flanked" (adjusted body posture) to maximize absorption of solar radiation on cool mornings; (iii) sought shade at midday to avoid adverse high temperatures. | (i) and (ii) raised body temperatures above ambient levels in the morning and maximized the amount of time that they could engage in feeding and reproduction before needing to perform (iii) to minimize heating later on. | Whitman (1988) |
| Endothermy | *Deilephila nerii*<br>(Lepidoptera: Sphingidae) | Moths beat and vibrated their wings pre-flight. | Warm up flight muscles to temperatures necessary to initiate flight. | Dorsett (1962) |
| Endothermy | *Vespa orientalis*<br>(Hymenoptera: Vespidae) | Hornet workers beat their wings at the nest entrance when temperature increased. | Ventilate the nest to prevent it from overheating. | Riabinin et al. (2004) |

### 4.2 Thermal orientation

In addition to seeking out temperatures that optimize their physiological functions, insects can use thermal information as an orientation cue, guiding them towards sources of food or mates. Similar to thermoregulation, there is good evidence that top-down integration of sensory information mediates thermal orientation behaviour. Antennal heat and cold receptors implicated in thermal orientation behaviour have been identified in several insects including mosquitoes (Davis and Sokolove 1975), triatomid bugs (Guerenstein and Lazzari 2009), and leaf-cutter ants (Ruchty et al. 2010a; Nagel and Kleineidam 2015). Scanning with antennal movements (Flores and Lazzari 1996), or thermoreception by specialized pit organs on the metathroax (Schmitz et al. 2007) allows these insects to detect minute temperature changes over short timescales, in the form of convective heat and/or infrared radiation (Davis and Sokolove 1975; Lazzari and Nunez 1989). In ants, information from antennal thermoreceptors is transmitted via thermosensitive neurons to glomeruli in the antennal lobe (Ruchty et al. 2010b), which may then project thermal information to higher brain centers involved in behavioural regulation, as in *Drosophila* thermoreception. Thermal orientation is used by blood-feeding insects to locate their hosts and initiate feeding, foraging ants to locate high-quality sun-exposed leaves, and fire-loving insects to locate sites for reproduction (see Table 5 for examples). For blood-feeding insects, temperature is part of a hierarchy of other cues used to locate hosts (e.g. host odours, carbon dioxide, moisture, visual information), and depends on temperature-sensitive aspects of physiology such as nutritional state (Guerenstein and Lazzari 2009; van Breugel et al. 2015). Thus, like thermoregulation, control of thermal orientation behaviour likely receives inputs from other sensory structures and is physiologically state-dependent (Fig. 2).

### 4.3 Thermosensory behavioural adjustments

Apart from behavioural thermoregulation and thermal orientation, do ectotherms use directly integrated thermal information to adaptively modify other aspects of their behaviour (Fig. 2) such as sex allocation, foraging strategies, mate choice, and social behaviour? To our knowledge, it has not yet been conclusively demonstrated that adaptive changes in such behaviours can arise from direct temperature perception. However, in several studies investigating the effects of temperature on insect behaviour, the way results are interpreted imply that high or low temperatures, or a sudden change in temperature, are being used as an indication of increased mortality risk (Dill et al. 1990), lower life expectancy (Moiroux et al. 2015), increased metabolic costs of activity (Traniello et al. 1994) or deteriorating environmental conditions (Dumont and McNeil 1992; Amat et al. 2006). A possible unstated premise behind these hypotheses is that the animals have the capacity to perceive temperature (as in thermoregulation), but use the integrated thermal information for other purposes. We will refer to these mostly hypothetical behavioural changes as Thermosensory Behavioural Adjustments. Although we present several possible examples here (see Table 6), their potential to fall into our category is provisional because, in several of these studies, the effect of temperature is difficult to tease apart from the effect of other variables (e.g. humidity, barometric pressure, photoperiod) (Dill et al. 1990; Dumont and McNeil 1992). Furthermore, in all of these previous studies, the observed changes in behaviour could have been due to temperature-induced metabolic/physiological constraints (short- or medium-term Kinetic effects) rather than Themosensory Behavioural Adjustments; an effort is not often made to distinguish between the two possibilities. In a rare effort to disentangle behavioural adjustments and physiological constraints, Moiroux et al. (2014) were able to measure ‘intended’ sex-ratio allocation and the corresponding secondary (realized) sex ratio by the haplodiploid egg parasitoid *Trichogramma euproctidis* (Girault) (Hymenoptera: Trichogrammatidae) at different temperatures by observing oviposition behaviour (pattern of abdominal contractions to indicate intended sex ratio) and rearing the Table 6. Possible examples of Thermosensory Behavioural Adjustments in insects, and their proposed functions.resulting offspring to adulthood. Similar to several other studies of parasitoid sex allocation (reviewed in King 1987), they observed a more male-biased secondary sex ratio at high and low temperatures compared to the intermediate temperature. However, the mechanisms behind the more male-biased sex ratios at the two extremes were different. At high temperatures, the intended proportion of male offspring (as well as the secondary sex ratio) increased, suggesting a temperature-dependent behavioural adjustment. At low temperature, the intended proportion of male offspring was the same as at intermediate temperature, but a reduction in the rate of egg fertilization (a Kinetic effect) caused a more male-biased secondary sex ratio than intended. Moiroux et al. (2014) hypothesized that intentionally increasing the proportion of males produced at higher temperatures is adaptive because developing and/or emerging at high temperatures may be less costly for male offspring than for female offspring. However, within our framework even this example can only be provisionally classed as a possible thermosensory behavioural adjustment, because it is not known whether temperature perception is actually involved, or rather whether the sex ratio adjustment is an indirect result of temperature's influence on another state variable (i.e., a medium-term Kinetic effect). Nonetheless, this work supports the notion that a single behavioural parameter can be subject to both compensatory adjustments and physiological constraints due to temperature, and illustrates the importance of distinguishing between the two possibilities when attempting to assign adaptive explanations to temperature-induced behavioural differences-in order for an adaptive explanation to be at all plausible, one first has to show that the behaviour is not simply due to a constraint.

**Table 5.** Examples of thermal orientation in insects, with proposed functions.

| Behavioural category | Species | Behaviour | Proposed function | Reference |
| --- | --- | --- | --- | --- |
| Foraging | *Atta vollenweideri*<br>(Hymenoptera:<br>Formicidae) | Worker ants were able to use thermal radiation as a learned orientation cue for food rewards. | Location and return to valuable resources such as sun-exposed leaves. | Kleineidam et al. (2007) |
| Foraging | *Triatoma infestans*<br>(Hemiptera: Reduviidae) | Bugs oriented towards heat sources that approximated the temperature of a food source. | Location of sites for blood-feeding. | Lazzari and Nunez (1989) |
| Foraging | *Melanophila acuminata*<br>(Coleoptera:<br>Buprestidae) | Beetles oriented towards thermal radiation emitted from forest fires. | Location of sites for mating and reproduction. | Evans (1964) |
| Foraging | *Aedes aegypti*<br>(Diptera: Culicidae) | Mosquitos oriented towards heat produced by their hosts, associating them with odor plumes using visual information. | Location of sites for blood-feeding. | van Breugel et al. (2015) |

**Table 6.** Possible examples of Thermosensory Behavioural Adjustments in insects, and their proposed functions.

| Behavioural category | Species | Behaviour | Proposed function | Reference |
| --- | --- | --- | --- | --- |
| Reproduction | *Trichogramma euproctidis*<br>(Hymenoptera: Trichogrammatidae) | Female parasitoids intentionally allocated more male offspring to hosts when foraging at high temperature. | Minimize fitness reduction of offspring caused by developing or emerging at high temperature (fitness of male offspring less affected). | Moiroux et al. (2014) |
| Foraging | *Aphidius ervi*<br>(Hymenoptera: Braconidae) | When at a high temperature, parasitoids more frequently attacked lower-quality host instars. | Adopting more risk-prone behaviours at high temperature was an adaptive response to a lower expectation of lifetime reproductive success due to a shorter life expectancy. | Moiroux et al. (2015) |
| Foraging | *Venturia canescens*<br>(Hymenoptera: Ichneumonidae) | Parasitoids spent longer on host patches following a drop in temperature. | Temperature drop used as indication of deteriorating conditions (increased cost of inter-patch travel). | Amat et al. (2006) |
| Dispersal | *Acyrtosiphon pisum*<br>(Hemiptera: Aphididae) | Aphids were less likely to drop off host plants when temperature was high and humidity was low. | Avoid leaving the host plant when probabilities of subsequent mortality are high. | Dill et al. (1990) |
| Mating | *Pseudaletia unipuncta*<br>(Lepidoptera: Noctuidae) | Males were less responsive to female sex pheromone (longer mating delay) when experiencing low-temperature, short day-length conditions. | Delaying mating under deteriorating end-of-season conditions provides a window for long-distance migration. | Dumont and McNeil (1992) |
| Communication | *Gryllus bimaculatus*<br>(Orthoptera: Gryllidae) | The responsiveness of female crickets to male songs at a given temperature was 'coupled' to the effect temperature has on song characteristics. | Permit mate recognition across range of variability in mating signals caused by temperature. | Doherty (1985) |

Perhaps the most thoroughly researched candidate thermosensory behavioural adjustment is “temperature coupling” (Gerhardt 1978) in animals that produce acoustic (e.g. crickets, grasshoppers) or visual (e.g. fireflies) signals for mate attraction. Several studies have observed that, across a range of sexually signaling species, levels of female responsiveness to male signals at a given temperature parallel temperature's effects on the male signal. That is, temperature's effect on male signal and the female's evaluation of the signal are coupled (reviewed in Greenfield and Medlock 2007). The adaptive explanation behind temperature coupling is that in order to properly assess male signals (i.e., permit species recognition and assess male quality), the evaluation of signals by females needs to be adjusted for the effect temperature has on male signals (Gerhardt 1978). Early on, two hypotheses were put forward to explain this phenomenon: the common neural elements hypothesis and the coevolution hypothesis (reviewed in Pires and Hoy 1992a). The common neural elements hypothesis proposes that male signal pattern generation and female song recognition are processed by homologous neural networks which are affected by temperature in the same way; for example, in crickets the female could use the temperature-dependent output from the thoracic Central Pattern Generator (CPG), which produces the neural signal used to generate stridulatory patterns in males, as a template for song recognition in the female brain. The coevolution hypothesis rather states that the neural networks underlying male pattern generation and female response are different, but have coevolved to sustain coupling of male signalling and female responsiveness over a range of temperatures. Despite more than 40 years of research and debate concerning the mechanisms behind temperature coupling (e.g., Gerhardt 1978; Pires and Hoy 1992a,Pires and Hoy 1992b; Doherty 1985; Bauer and von Helverson 1987; Shimizu and Barth 1996; Ritchie et al. 2001; Greenfield and Medlock 2007), its underlying mechanisms and adaptive significance remain unclear. In the context of our framework for temperature's effects on behaviour, several of the behaviour types we outline have been proposed as a mechanism under the umbrella of both of the above hypotheses. For example, a modified version of the common neural elements hypothesis proposed by Pires and Hoy (1992b) states that female crickets integrate temperature-dependent input from the thoracic CPG in the brain to produce responses to male signals (short- or medium-term Kinetic effect), possibly with input from thermosensors in the head or the thorax (in which case it would be classed as a Thermosensory Behavioural Adjustment). In both of these cases, it is unclear if signal generation in males would only be due to temperature's effect on metabolic processes and speed of nerve signal transmission in the CPG (Indirect Kinetic effect) or less plausibly, if CPG signals would also be integrated with thermosensory input in males' brains to adjust stridulatory patterns (Thermosensory Behavioural Adjustment). Greenfield and Medlock (2007) presented evidence that temperature coupling in the acoustically-mating moth *Achroia grisella* (F.) (Lepidoptera: Pyralidae) does not even constitute a true adaptation, and the apparent coordination of signal generation and reception simply represents an emergent effect of the common influence of temperature on the underlying neuromuscular physiology of males and females (short- and medium-term Kinetic effects), supporting the original common neural elements hypothesis. Clearly, further investigation concerning the physiological mechanisms behind temperature coupling are still needed.

Despite the current lack of clear evidence of Thermosensory Behavioural Adjustments, one could speculate under what conditions we would expect thermal information to be used to adjust behavioural strategies rather than simply moving to a different temperature. Given the inertial evolutionary effect of thermoregulatory behaviours (Huey et al. 2003), we might predict that compensatory behavioural adjustments could evolve when the costs of behavioural thermoregulation are too high or thermoregulatory behaviours are not feasible. This could occur, for example: (i) when opportunities for reproduction or feeding are in a fixed location and the location of alternative opportunities in better thermal environments is impossible or suboptimal, so that continuing to exploit the resource necessitates behaviourally compensating for the imposed thermal environment; (ii) when temperature's effect is external to the individual and cannot be changed by thermoregulatory means (e.g. in female crickets intercepting a male courtship call determined by his body temperature); or, (iii) when options for thermoregulation by microhabitat selection are extremely limited by, for example, very poor dispersal ability (e.g. in immature stages of some insects) or close associations with other species (e.g. plant-herbivore or host-parasitoid) (Rand and Tscharntke 2007; Tylianakis et al. 2008).

## 5. Using the framework in thermal behavioural ecology

In building our framework for the effect of temperature on ectothermic animal behaviour, we have encountered two major themes. First, apart from thermoregulatory behaviours, there have been very few tests of whether temperature influences adaptive decision-making processes in insects. Second, even when an apparently adaptive behavioural modification is observed with changing temperature, it is unknown whether it is a direct response to temperature perception (Integrated effect) or rather just a medium- or long-term Kinetic effect of temperature on some aspect of the animal's physiological state. To further complicate the matter, all the different effects we have synthesized here are often acting simultaneously in natural populations, the behaviour of an individual insect observed at any given point of time is likely to be influenced by the temperature at which it developed (long-term Kinetic effects), the temperature it experienced the previous days and hours (medium-term Kinetic effects), the current ambient temperature (short-term Kinetic effects), as well as intentional behavioural control due to top-down integration of thermal information (Integrated effects). Furthermore, as previously discussed in the context of thermoregulation, the effects of integrated thermal information on behaviour may be dependent on aspects of physiological state that are influenced by on short-, medium- and long-term Kinetic effects of temperature (Fig. 2).

To appreciate how Kinetic and Integrated effects can interact to produce a behavioural phenotype, it is helpful to consider changes in insect walking patterns across a hypothetical temperature gradient, considering only short-term physiological (Kinetic) effects and thermoregulation (an Integrated effect). Insect walking is controlled by clusters of neurons termed central pattern generators (CPGs) in the central or peripheral nervous system, which control the firing of motor neurons that activate the muscles responsible for limb movement (Delcomyn 2004; Büschges et al. 2008). The signals received by the motor neurons and the resulting muscle movements are further modulated by sensory feedback (e.g. from visual systems, proprioreceptors, mechanosensory organs, hygro- and thermo-sensory organs), resulting in changes of the characteristics of walking behaviour such as direction and speed. When the insect walks across the temperature gradient, observed behavioural modifications due to temperature would have both Kinetic and Integrated components. In terms of Kinetic effects, changing temperature would result in changed neuronal conduction speed, a different firing rate of CPG neurons, a change in the rapidity of neurotransmission at neuromuscular junctions, and modified metabolic rates in muscle tissue due to differences in available kinetic energy (Robertson and Money 2012); all of which would result in the insect tending to walk faster at higher temperatures and slower at lower temperatures. Simultaneously, Integrated effects would also affect walking behaviour: the animal would receive feedback regarding temperature changes from thermosensory organs, the thermal information would be encoded by the brain and used to modulate muscular output; this in turn would bias walking towards the individual's ‘preferred’ temperature. Thus, Integrated and Kinetic effects occur simultaneously to produce the observed behavioural output, but teasing apart these two kinds of effects has seldom been attempted.

To help guide future research in this area, we suggest two general, complementary approaches to distinguish integrated effects from direct and indirect kinetic effects: behavioural kinetic null modelling and behavioural ecology experiments using temperature-insensitive mutants. In both cases, the principle is the same: *if one can demonstrate that a behavioural modification does not occur because of direct temperature perception, it is mostly likely due to a combination of Kinetic Effects rather than an Integrated effect.*

### 5.1 Behavioural Kinetic null modeling

To determine whether a change in an animal's behaviour is a direct response to temperature perception, one can ask: how would the organism behave in the absence of Integrated effects? This kinetic null model would contain relationships describing only Kinetic effects of temperature on behaviour (i.e., constraints), ranging from short-term to long-term. Thus, the null model would treat the animal as a “metabolic machine” whose behaviours are driven only by the bottom-up Kinetic effects of temperature on the rates of biochemical processes. Then, one or more Integrated models could be constructed assuming that the animal's behaviour is optimized to the impacts of temperature on fitness payoffs; that is, the individual is monitoring temperature and intentionally modifying its behaviour to suit thermal conditions (i.e., there are likely Integrated effects acting on behaviour).

Null models have been used previously in the context of behavioural thermoregulation. Huey et al. (2003) considered how the body temperature and sprint performance of thermoregulating lizards would vary over an elevational gradient, compared to “null lizards” that did not perform any thermoregulatory behaviours. Because of thermoregulation, the empirically measured body temperatures of real lizards declined less rapidly with increasing elevation than would be expected of null lizards. Vickers and Schwarzkopf (2016) added spatial and temporal components to a null model of behavioural thermoregulation. This new model also estimates the fitness costs associated with imprecise thermoregulation, and introduces a metric to assess what the authors term “effort”, constituting deliberate behavioural input into thermoregulation (and hinting at what we call Integrated Effects). Other studies, using the nematode *Caenorhabditis elegans* (Anderson et al. 2007) and *Drosophila* (Dillon et al. 2012) as model organisms, formulated null models in order to disentangle temperature's effects on movement from temperature preference on the spatial distribution of small ectotherms across a thermal gradient. Here, the null models would assume random (Brownian) motion whose rate is dependent on temperature's short-term Kinetic effects on movement. These studies showed that the expected null distributions were not necessarily intuitive and were extremely dependent on experimental conditions (i.e., the range of temperatures on the gradient relative to the organism's thermal tolerance). Thus, these null models are critical to establish what proportion of the observed distribution of small ectothermic species on a thermal gradient depends on thermal preference (Integrated effects) and not Kinetic effects. So, while Kinetic null modelling is beginning to be fruitfully applied to assessing fitness benefits of thermoregulation and teasing out the intentional/deliberate component of thermoregulatory behaviours, it has not yet been used to investigate potentially adaptive components of other behavioural responses to temperature.

As an example of how the kinetic null modelling approach could be extended to other complex thermally dependent behaviours, consider a time-limited forager in an environment where resources are patchily distributed - a classic behavioural ecology paradigm. Optimal foraging theory postulates that such a forager should adjust the amount of time spent in each patch (its Patch Residence Time; PRT) based on factors (e.g. internal physiological state, relative host or prey quality in each patch, travel time between patches, stochastic mortality risk) that influence the forager's perception of future fitness gains (reviewed in Wajnberg 2006). A forager is predicted to leave a resource patch when its rate of fitness gain on the patch falls below the average rate of fitness gain in the environment (Charnov 1976). When PRT varies with temperature, potential explanations may include both Kinetic and Integrated effects. Kinetic effects may include increased movement speed, more rapid host or prey handling, higher digestion rate (herbivores and predators), and more rapid oogenesis (parasitoids). The thermal dependency of such parameters could be measured and used to parameterize a null model for the effect of temperature on PRT. Other parameters would have fixed values whose thermal dependencies are unknown to the forager, because they do not depend on some aspect of its own metabolic rate. Then, Integrated models could be built assuming that the forager's behaviour is adapted to the thermal dependency of one or more of these other factors (e.g. travel time, host quality, mortality risk, energy expenditure) affecting the trade-off between current and future fitness gains. One would then test the effect of temperature on forager PRT and determine if the data best fit the predictions of the Kinetic null model or one of the alternative Integrated models. If the data do not fit any of the models, then there are two possibilities: (i) there are relevant Kinetic effects (e.g. medium- or long-term) that are not being included in the null model, or (ii) the hypotheses underlying the Integrated models are flawed. The models could then be updated with new hypotheses concerning how the behaviour is affected by temperature and tested with new experiments, gradually eliminating competing hypotheses and improving the fit between the model and experimental data. Used judiciously, this approach could be extended to a wide range of behaviours including foraging (e.g. resource choice, resource defense), reproduction (clutch size, mate guarding duration), communication (calling behaviour, vibrational signalling), dispersal (selection or time allocation amongst different habitats), and anti-predator strategies (choice to attack or flee).

This proposed null modelling approach has some limitations. Firstly, parameterizing a useful Integrated model requires an accurate knowledge of how temperature affects the fitness payoffs of different possible behavioural decisions and assumes that there have not been evolutionary constraints on the animal's ability to adapt to these differential payoffs. Thus, as with all models, these Integrated models will only be as good as the hypotheses contained within them. Second, to include all relevant Kinetic effects in a null model, one needs a complete understanding of how thermal physiology underlies behaviour to make sure that deviations from the null model reflect Integrated effects rather than Indirect Kinetic effects that have not been accounted for. This should sometimes be apparent by a poor fit of the experimental data to the Integrated models, although it is conceivable that an Indirect Kinetic effect could produce the same behavioural pattern as if the animal were directly perceiving temperature (for example, if an individual is “tracking” its own metabolic rate). So, in some cases, it may also be necessary to employ the next approach, the use of temperature-insensitive mutants.

### 5.2 Use of TEmperature-Insensitive Mutants In Behavioural Ecology Experiments

The most direct way to test if direct temperature perception is involved in a behaviour would be to examine if individuals with dysfunctional temperature sensation still change their behaviour over a range of temperatures (i.e., when Integrated effects are “silenced”). So far, this has only been performed in the context of *Drosophila* behavioural thermoregulation (see beginning of section 4). Temperature-insensitive *Drosophila* have been produced either by developing mutant lines deficient in thermoreceptor function (Hamada et al. 2008; Gallio et al. 2011; Shen et al. 2011), functionally inactivating cellular structures involved in temperature perception with transgenic expression of toxins (Gallio et al. 2011), or the removal/ablation (e.g. with lasers) of putative thermoreceptory organs (Sayeed and Benzer 1996; Hamada et al. 2008). While these studies have been made possible by the wide array of genetic tools available for *Drosophila*, the recent sequencing of many other insect genomes and the recent advent of greatly improved gene-editing technology (Kistler et al. 2015) could facilitate the extension of similar tools to other biological systems. These methodologies, alone or in combination, could be used to develop lines of temperature-insensitive mutants of other insect species that could then be genetically characterized and tested for temperature-dependent behavioural adjustments. Of course, this proposed approach assumes that the thermoreceptory mechanisms involved in thermoregulation, including the location and function of specific thermosensory organs, are the same as those involved in Thermosensory Behavioural Adjustments. This matter could be subjected to preliminary investigation by, for example, conducting investigations of temperature coupling in temperature-insensitive mutant lines of *Drosophila* (Ritchie et al. 2001). Insect parasitoids are another group of potentially ideal biological models for testing this approach. Using parasitoids, one could test whether temperature-dependent sex ratio allocation (Moiroux et al. 2014) or patch residence time adjustments with temperature (Amat et al. 2007) depend directly on temperature perception by comparing responses of mutant versus wild-type lines, or using gene editing to knock out genes known to be critical for temperature perception. This methodology could be particularly effective if used in combination with null modelling-one would expect the behaviour of temperature-insensitive individuals to follow the predictions of the null model, while the behaviour of wild-type individuals should follow the predictions of the integrated model.

## 6. Conclusions

To be able to predict the global effects of temperature changes on ecosystems, a detailed understanding of how it acts on all levels of biological organization is needed. When investigating the influence of temperature and the potential effects of global warming on communities and ecosystems, it is typical to consider mainly physiology-based changes in life histories of organisms such as changes in phenology (for a review, see Parmesan 2006). However, the influence of temperature on behaviours may sometimes explain ecological changes associated with global warming better than its influence on metabolism or physiology alone (e.g., Barton 2010, Harmon and Barton 2013). As we outlined in this paper, behavioural responses to temperature can result from physiological constraints, adaptive behavioural changes due to temperature, and often both. Distinguishing among these causes is important, because different underlying mechanisms may have different capacities to adapt to changing environments. The framework we have presented here constitutes a first attempt to organize our current understanding and focus future research on the role temperature plays in behavioural ecology. Researchers planning to investigate how ectothermic animal behaviour changes with temperature can use this framework as a way to think about how organisms *should* change their behaviour with temperature, and how constraints will simultaneously act to masquerade as, or obscure, adaptive components of the behavioural response. As understanding advances about how thermal physiology underlies behaviour, our framework needs to be modified and built upon. Hopefully in the meantime it will help guide more focused, hypothesis-based research, and lead to a fuller consideration of temperature's effects on the behavioural ecology of ectothermic animals.

## Acknowledgements

We thank Mike Angilletta and Kevin Tougeron for providing insightful comments on an earlier draft of the manuscript, and Alexandre Leblanc and Jean-Philippe Parent for helpful discussions. This work was supported by a Natural Sciences and Engineering Council of Canada Postgraduate Scholarship to P.KA. and the Canada Research Chair in Biological Control to J.B.

